# Oral Regeneration Is the Default Pathway Triggered by Injury in *Hydra*

**DOI:** 10.1101/2020.07.06.189811

**Authors:** Jack Cazet, Celina Juliano

## Abstract

In animals capable of whole-body regeneration, a single bisection injury can trigger two different types of regeneration. Currently, it is not well understood how this adaptive response is transcriptionally regulated. We therefore comprehensively characterized transcript abundance and chromatin accessibility during two types of regeneration—oral and aboral regeneration—in the cnidarian *Hydra vulgaris*. We found that there was an initial generalized response to injury that was not dependent on the type of structure being regenerated. Canonical Wnt signaling, which specifies oral tissue in cnidarians, was activated during this generalized response, likely through the direct upregulation of Wnt ligands by conserved injury-responsive transcription factors. As regeneration progressed, transcription diverged and Wnt signaling became restricted to oral regeneration. We found that TCF, the transcription factor downstream of Wnt signaling, was required for the initiation of oral and aboral-specific transcription, suggesting that the Wnt pathway is important for both types of regeneration. Finally, we found that Wnt signaling was also activated by puncture wounds and that removing pre-existing organizers induced these injuries to undergo ectopic head regeneration. Our work suggests that injuries activate an oral regeneration program, and that other regeneration outcomes are caused by signals from the surrounding tissue. Furthermore, these findings suggest that Wnt signaling may be part of an ancient and conserved metazoan wound response program predating the split of cnidarians and bilaterians.

## Introduction

Regeneration is an injury-induced morphogenetic process that enables the restoration of lost or damaged body parts. Although nearly all animals are capable of some form of regeneration, the greatest regenerative capacity is found in the invertebrate species capable of rebuilding their entire body from small tissue fragments through a process called whole-body regeneration. When these animals are bisected, regeneration in the two resulting tissue fragments gives rise to two morphologically identical individuals. Importantly, if the bisection does not fall along a plane of symmetry (i.e. the amputation transects a body axis), the structures regenerated by each tissue fragment are different. This raises the question of how a single injury gives rise to two different types of regeneration.

In diverse animal regeneration models, the fate of regenerating tissue is directed by conserved developmental patterning pathways—such as Wnt and BMP signaling—that are activated following injury (Gemberling et al., 2013; Kawakami et al., 2006; Owlarn and Bartscherer, 2016; Srivastava et al., 2014). However, it is not well understood how these pathways are activated by injury, nor is it well understood how their activation is spatially or temporally restricted to ensure the restoration of the original body plan. In particular, the gene regulatory networks (GRNs) responsible for coordinating the transcriptional response during regeneration, including the activation and modulation of patterning pathways, are not well characterized in most regeneration models.

*Hydra vulgaris* is a freshwater cnidarian polyp with an exceptional regenerative capacity, capable of fully rebuilding its body from tissue pieces containing as few as 300 cells, or ∼1% of an adult polyp (Shimizu et al., 1993). *Hydra* has a simple body plan defined by a single oral-aboral axis, with tentacles and a hypostome at the oral end, referred to as the head, and adhesive cells at the aboral end, referred to as the foot (Fig. 1A). In intact *Hydra*, tissue polarity is determined by two stable self-maintaining organizers, a head organizer and a foot organizer, that are found at the poles of the oral-aboral axis. Importantly, regenerating *Hydra* that have lost one or both of these organizers maintain their polarity. Thus, an oral-facing wound will regenerate a head and an aboral-facing wound will regenerate a foot (Webster, 1971) (Fig. 1B). This suggests that the surrounding uninjured tissue—referred to as the injury’s tissue context—provides cues that direct positional specification during regeneration. While regeneration in *Hydra* is typically completed within ∼3 days, the organizers that dictate the positional identity of the regenerating tissue generally form within 12 hours post-amputation (hpa) during a process called pre-patterning (J. Hicklin and Wolpert, 1973; MacWilliams, 1983a; Petersen et al., 2015). Information from the surrounding tissue context is therefore incorporated into the injury response relatively quickly, but the mechanism underlying this process is unknown.

**Figure 1.**
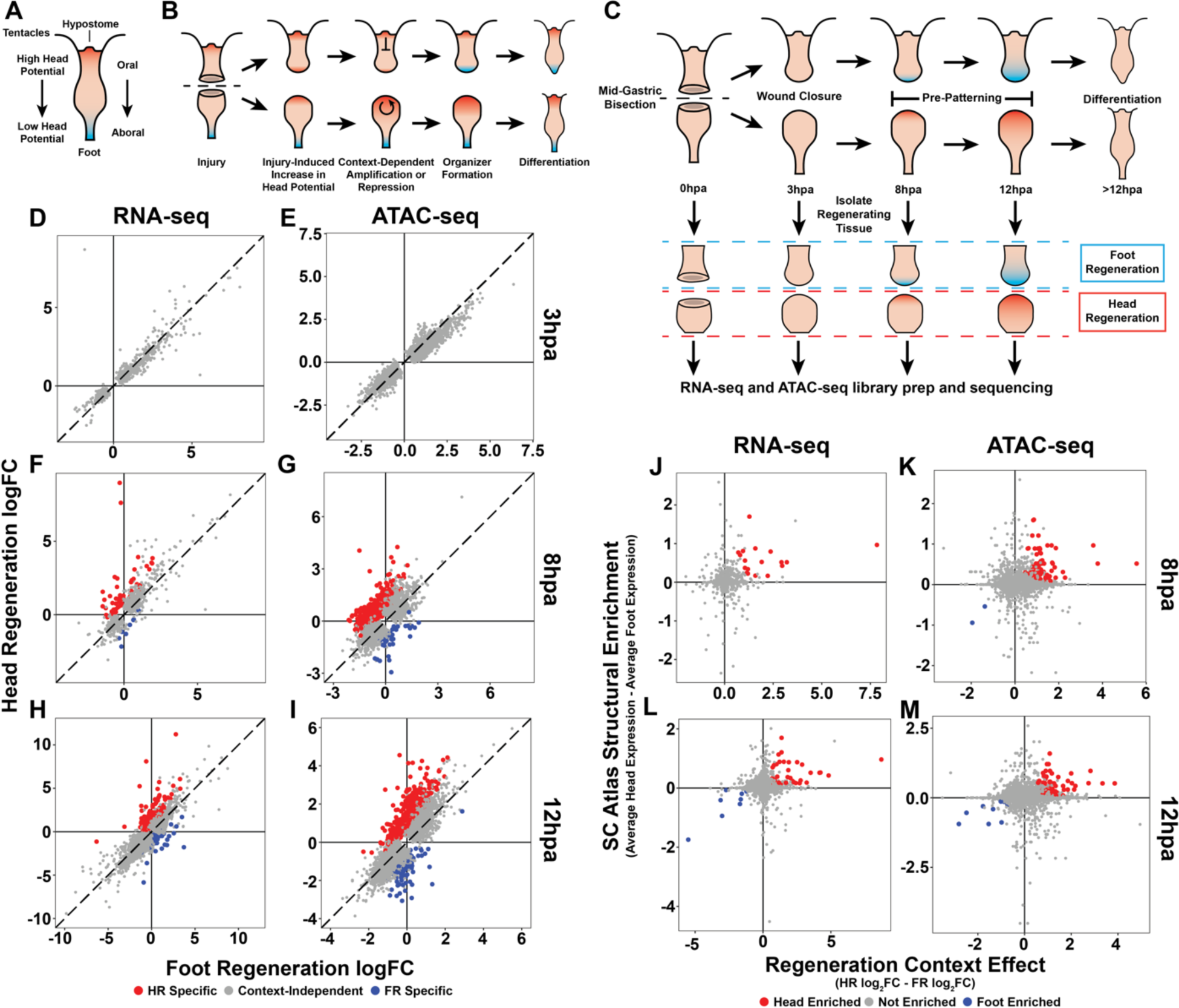
The initial transcriptional response to mid-gastric bisection is context independent but diverges by 8 hpa. (A) A diagram of the *Hydra* body plan. Red coloration indicates a high capacity to form a new head organizer. Blue coloration indicates a low capacity to form a new head organizer. (B) A diagram of the oral induction model. (C) Phases of regeneration and the experimental design for ATAC-seq and RNA-seq library generation. (D-I) Comparison of average log_2_ fold change (log_2_FC) in transcript abundance (D,F,H) or chromatin accessibility (E,G,I) during head and foot regeneration at 3, 8, and 12 hpa. The transcriptional response during head and foot regeneration is identical at 3 hours post-amputation (hpa) but becomes distinct by 8hpa. Features that showed significant differences in the injury response during head and foot regeneration when compared to 0 hpa controls are highlighted in red for head regeneration-specific features or blue for foot regeneration-specific features (FDR ≤ 1e-3 for RNA-seq and FDR ≤ 1e-4 for ATAC-seq). The dotted line indicates perfect correlation between regeneration types. (J-M) Plots comparing enrichment at the oral/aboral poles in uninjured *Hydra* and during regeneration. Genes associated with uninjured head tissue were specifically upregulated in regenerating heads by 8 hpa. Genes associated with uninjured foot tissue were specifically upregulated in regenerating feet by 12 hpa. Enrichment in uninjured head and foot tissue (i.e. SC Atlas Structural Enrichment) was determined by comparing the average relative expression levels in epithelial head cells isolated from the *Hydra* single cell atlas to expression in epithelial foot cells. A positive fold change indicates enrichment in head tissue and a negative fold change indicates enrichment in foot tissue. Head or foot specific activation during regeneration (i.e. Regeneration Context Effect) was calculated by subtracting the log_2_FC in regenerating head tissue from the log_2_FC in regenerating foot tissue using the 0 hpa timepoint as the control. A positive fold change indicates higher activation during head regeneration and a negative fold change indicates higher activation during foot regeneration. ATAC-seq peaks (J,L) or transcripts (I,K) that were activated in a head regeneration-specific manner that were also enriched in uninjured head tissue are highlighted in red. ATAC-seq peaks or transcripts that were activated in a foot regeneration-specific manner that were also enriched in uninjured foot tissue are highlighted in blue.

Early work studying patterning during *Hydra* regeneration led to the hypothesis that positional identity in regenerating tissue is determined by the balance between a head-promoting signal generated by injured tissue and a diffusible head-inhibiting signal produced by head and head-proximal tissue (Gierer and Meinhardt, 1972; Kobatake and Sugiyama, 1989; MacWilliams, 1983a, 1983b). This hypothesis, which we have termed the oral induction model, proposed that when the level of inhibition is sufficiently low—such as when the head has been removed—the head promoting signal from the injury initiates a feed-forward loop that induces the formation of a new self-maintaining head organizer. However, during foot regeneration, it was hypothesized that the inhibition originating from oral tissue prevents the initial injury signal from triggering the feed-forward loop, causing head organizer formation to fail and resulting instead in foot specification (Fig. 1B). Although this model successfully accounts for a number of documented patterning phenomena, it has not been tested using molecular tools. Consequently, the molecular mechanisms underlying the activation of an oral patterning cascade following injury and the subsequent modulation of that cascade by the surrounding tissue context remain largely unknown.

Over a decade after the oral induction model was proposed, canonical Wnt signaling was discovered as the primary regulator of oral-aboral positional identity in *Hydra*. Canonical Wnt ligands are expressed in homeostatic and regenerating heads (Hobmayer et al., 2000; Lengfeld et al., 2009), and Wnt signaling is both necessary and sufficient to induce the formation of ectopic head-like structures (Broun et al., 2005; Gee et al., 2010; Gufler et al., 2018). However, Wnt signaling dynamics during wound repair have been largely unexplored outside the context of head regeneration. Furthermore, although recent research has begun to illuminate the transcriptional changes that occur as regenerating *Hydra* tissue undergoes positional specification (Gufler et al., 2018; Petersen et al., 2015; Wenger et al., 2019), the regulatory mechanisms by which Wnt signaling and other pathways are activated in a context-specific manner are not well understood.

We therefore sought to better understand the molecular regulatory logic directing positional specification during regeneration in *Hydra*. To this end, we profiled transcript abundance and chromatin accessibility genome-wide during the first 12 hours of head and foot regeneration following mid-gastric bisection. We found that the early transcriptional response to injury was virtually identical during head and foot regeneration and included the activation of canonical Wnt signaling, which we propose is directly triggered by injury responsive basic leucine zipper (bZIP) transcription factors (TFs). By 8 hpa, we found clear evidence of transcriptional divergence between head and foot regeneration, which coincided with Wnt signaling becoming restricted to head regeneration. In addition, we found that the Wnt-responsive TF TCF was required for the global initiation of pre-patterning and transcriptional divergence during both head and foot regeneration, suggesting TCF has other roles in addition to head specification. Finally, we found that non-amputation injuries also activated Wnt signaling and could induce ectopic head regeneration when pre-existing organizers were removed. Overall, our findings demonstrate that injuries trigger an oral patterning cascade regardless of tissue context through the activation of canonical Wnt signaling, and that the outcome of this cascade is regulated by long range signals generated by pre-existing organizers. Our work therefore strongly supports the oral induction model and provides insight into the regulatory logic directing whole-body regeneration in *Hydra*.

## Results

### The transcriptional response to injury is initially independent of tissue context but becomes specialized between 3 and 8 hours post-amputation

To better understand how regenerating tissue acquires the appropriate positional identity in *Hydra*, we generated RNA-seq and ATAC-seq (Buenrostro et al., 2013; Corces et al., 2017) libraries from polyps undergoing either head or foot regeneration at 0, 3, 8, and 12 hours after mid-gastric bisection (Fig. 1C; Supplementary Tables 1 & 2). ATAC-seq enables the identification of active cis-regulatory elements genome-wide by generating high-throughput sequencing libraries that are highly enriched for accessible regions of chromatin. Because mid-gastric bisection triggers two different types of regeneration from the same injury, these data allowed us to characterize how differences in tissue context affected cis-regulatory element activity and transcriptional output during early regeneration. We used mid-gastric bisection to minimize positional biases, as amputations closer to the extremities will more rapidly regenerate the structure they are closest to (J. Hicklin and Wolpert, 1973; Webster, 1971). We chose 12hpa as our latest timepoint because previous research demonstrated that new organizers are established by approximately this time (J. Hicklin and Wolpert, 1973; MacWilliams, 1983a).

We first sought to determine which aspects of the transcriptional response to injury were influenced by tissue context. We accomplished this by identifying the Injury <u>R</u>esponsive ATAC-seq <u>P</u>eaks (IRPs) or Injury <u>R</u>esponsive <u>T</u>ranscripts (IRTs) that changed in significantly different ways during the two types of regeneration (Supplementary Files 1 and 2). At 3 hours post amputation (3 hpa), we found that the changes in transcript abundance and chromatin accessibility during head and foot regeneration were indistinguishable (Fig. 1D,E). Our data therefore suggest that the initial transcriptional response to amputation is not influenced by tissue context. This finding is consistent with previous research in both planarians and cnidarians that documented an initial context-independent stage of regeneration (Schaffer et al., 2016; Wenger et al., 2019; Wurtzel et al., 2015).

At 8 hpa, however, we found clear evidence of context-dependent transcription, with 63 IRTs and 464 IRPs showing significant differences when comparing head and foot regeneration (Fig. 1F,G). We postulate that the greater number of differentially activated IRPs relative to IRTs is due to gene loci often being associated with multiple peaks. In addition, certain cis-regulatory elements may not be sufficient to meaningfully change transcript abundance. Transcriptional differences continued to accumulate as regeneration progressed, with 139 IRTs and 631 IRPs exhibiting context-dependent differences at 12 hpa (Fig. 1H,I). Our analysis therefore suggests that tissue context is integrated into the transcriptional response to amputation between 3 and 8 hpa, which is consistent with a recent transcriptomic study of *Hydra* head and foot regeneration that estimated the time of divergence to be between 4 and 8 hpa (Wenger et al., 2019).

Importantly, we found that the context-dependent IRTs that we identified were significantly enriched in nearby context-dependent IRPs (14.1-fold enrichment at 8 hpa, P=3.6e-25; 9.8-fold enrichment at 12 hpa, P=3.7e-40), demonstrating a meaningful correspondence between our ATAC-seq and RNA-seq datasets. Furthermore, we found that there was a significant overlap in the context-dependent IRTs identified in our study when compared to the context-dependent IRTs identified from comparable timepoints in a previously released RNA-seq dataset (507-fold enrichment at 8hpa, P∼0; 35-fold enrichment at 12hpa, P=1.11e-93) (Wenger et al., 2019). We therefore found that context-dependent transcription during regeneration was highly reproducible across independent studies and across orthogonal high-throughput sequencing methodologies.

### Wnt signaling initially increases during both head and foot regeneration but becomes head specific by 8 hpa

For regeneration to restore head and foot tissue, part of the injury response must include the re-activation of the pathways that specify and maintain these structures under steady-state conditions. We therefore looked for evidence of the re-activation of head or foot specific pathways during regeneration to identify the genes involved in the early stages of new head or foot organizer formation. To do this, we focused on the earliest transcripts to exhibit context-dependent transcription during regeneration and characterized their expression profiles in uninjured *Hydra* using a recently published single cell sequencing atlas (Siebert et al., 2019).

At 8 hpa, the earliest time point in our dataset with transcriptional differences between head and foot regeneration, there were 55 IRTs and 426 IRPs that showed significantly higher activation during head regeneration relative to foot regeneration. We found that 23/55 of the head regeneration-specific IRTs and 65/426 of the head regeneration-specific IRPs were also significantly enriched in uninjured head tissue (Fig. 1J,K). These early head-specific factors included previously characterized genes associated with the *Hydra* head such as *wnt9/10c* (Lengfeld et al., 2009), *wnt3* (Hobmayer et al., 2000), *brachyury1* (Technau and Bode, 1999), *naked cuticle* (Petersen et al., 2015), and *sp5* (Vogg et al., 2019) (The *Hydra* 2.0 genome gene model IDs for the gene names used throughout this study can be found in Supplementary Table 3). In contrast, we did not observe a similar activation of foot-specific transcription at 8 hpa, although there was some evidence of foot specific chromatin remodeling (Fig. 1J,K). At 12 hpa, however, several genes enriched in homeostatic foot tissue, including the known foot-associated TFs *distal-less* (Hemmrich et al., 2012) and *nk-2* (Grens et al., 1996) (Fig. 1L,M), were upregulated in a foot regeneration-specific manner. We hypothesize that these early context-specific genes reside at the top of the regulatory hierarchy directing the specification of head or foot tissue during regeneration.

We noted that 47/101 of the transcripts that were specific to head regeneration by 12 hpa were initially upregulated during both head and foot regeneration at 3 hpa (Fig. 2A). These transcripts therefore exhibited both context-independent and context-dependent phases of expression. Notably, these biphasic IRTs included the Wnt signaling ligands *wnt9/10c* and *wnt3* and the Wnt secretion facilitator *wntless* (Fig. 2B-D). This motivated us to determine if other Wnt pathway components showed similar expression dynamics. We used KEGG pathway annotations to systematically identify and characterize the expression of Wnt signaling pathway components in uninjured *Hydra* and during regeneration (Supplemental Fig. 1A,B). We found that the head-specific Wnt signaling-associated genes *wnt7, dishevelled* and *sp5* were significantly upregulated at 3 hpa during foot regeneration. Although the head regeneration differentials for these genes did not pass our significance cutoff at 3 hpa, all three transcripts were significantly upregulated during head regeneration by 8hpa (Fig. 2E-F, Supplemental Fig. 1B). These findings raised the possibility that Wnt signaling, which is crucial for specifying oral tissue during *Hydra* regeneration, is initially activated as part of a rapid context-independent response to injury.

**Figure 2.**
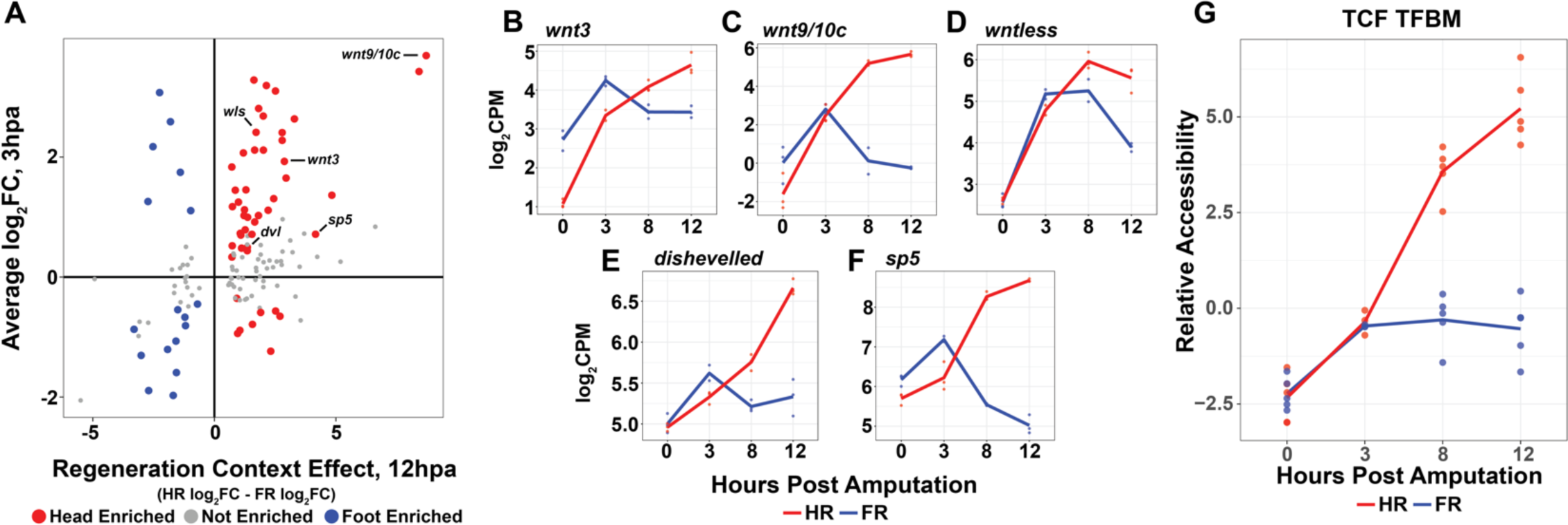
Wnt signaling is initially activated in a structure independent manner but becomes restricted to regenerating head tissue by 8 hpa. (A) Comparison of context-dependent transcription at 12hpa and the log_2_FC at 3hpa relative to 0hpa. Head and foot regeneration-specific transcripts are initially activated in a context-independent manner at 3hpa. Transcripts that were differentially expressed from 0 to 3 hpa and were activated in a head or foot regeneration-specific manner at 12hpa are highlighted in red and blue respectively. Log_2_FC at 3hpa was calculated as the average log_2_FC across both head and foot regeneration. (B-F) RNA expression plots showing average normalized RNA-seq read counts for Wnt signaling components in log_2_ counts per million (log_2_CPM) during head and foot regeneration. Wnt-signaling components are initially upregulated during head and foot regeneration but become head regeneration-specific by 12hpa. (G) Average relative chromatin accessibility plot of peaks containing the TCF transcription factor binding motif (TFBM) during head and foot regeneration. The TCF TFBM is associated with increases in accessibility during both head and foot regeneration at 3hpa, but later increases in accessibility are restricted to head regeneration. HR: head regeneration, FR: foot regeneration.

Although canonical Wnt signaling components were upregulated during both head and foot regeneration, it was not clear if this resulted in the downstream activation of Wnt responsive transcription. We therefore evaluated Wnt signaling activity during regeneration by using chromVAR (Schep et al., 2017) to quantify the average change in chromatin accessibility near predicted binding sites of TCF, the TF downstream of canonical Wnt signaling. We found that chromatin accessibility near TCF transcription factor binding motifs (TFBMs) was correlated with injury-responsive Wnt ligand expression during regeneration, with accessibility initially increasing during both head and foot regeneration from 0 to 3 hpa but then increasing in a head-specific manner from 8 hpa onward (Fig. 2G). These data therefore suggest that the Wnt pathway is initially activated by injury in a structure-independent manner but then becomes restricted to head regeneration as information from the surrounding tissue is integrated into the transcriptional response. This observation is consistent with the oral induction model, which states that injuries initially promote the formation of a new oral organizer regardless of how the injury is ultimately resolved.

### TCF inhibition delays the onset of tissue context-dependent transcription during head and foot regeneration

Our results suggested that amputation caused an increase in Wnt signaling during both head and foot regeneration from 0 to 3 hpa. Although the function of Wnt signaling during head regeneration is well established, possible roles for Wnt signaling during foot regeneration were unclear. Recent research found that the small molecule TCF inhibitor iCRT14 (Gonsalves et al., 2011) blocked both head and foot regeneration in *Hydra*, suggesting that the transient pulse of Wnt signaling we observed could be necessary for foot regeneration (Gufler et al., 2018). We therefore sought to better understand the role of canonical Wnt signaling during head and foot regeneration on a genome-wide scale by treating regenerating *Hydra* with 5µM iCRT14 and generating RNA-seq and ATAC-seq libraries at 0, 8, and 12 hpa after mid-gastric bisection. We additionally collected a 3 hpa timepoint for our iCRT14-treated ATAC-seq dataset.

We first assessed iCRT14’s effects on the chromatin accessibility near TCF TFBMs to validate that the drug treatment successfully inhibited TCF. As expected, we observed a diminished and delayed increase in average TCF TFBM accessibility during head regeneration (Fig. 3C). Consistently, the upregulation of numerous head regeneration specific genes—including *axin, prdl-a, sFRP, tcf, gremlin-like*, and *budhead*—was significantly reduced or blocked altogether in iCRT14-treated regenerates at 8 hpa (Fig. 3A). These data therefore indicate that iCRT14 treatment attenuated TCF-dependent transcriptional activation during regeneration.

**Figure 3.**
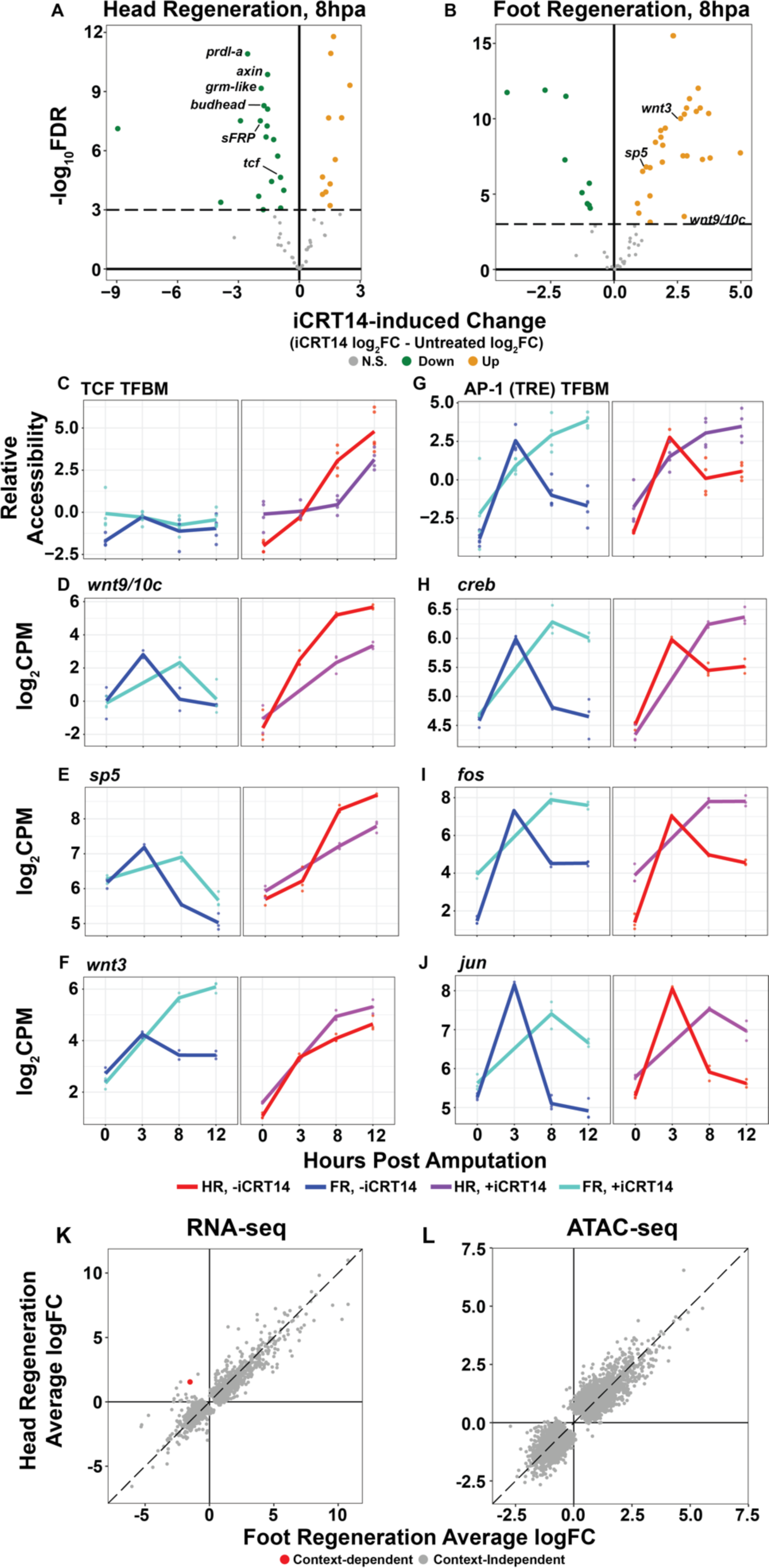
TCF is required for the initiation of repatterning during head and foot regeneration. (A,B) Plots depicting the magnitude (i.e. log_2_FC) and significance (i.e. -log10(FDR)) of the iCRT14-induced changes in the transcriptional response to injury at 8 hpa for (A) head and (B) foot regeneration. iCRT14 significantly attenuated the activation of head specific transcription during head regeneration and delayed the context-specific downregulation of Wnt signaling transcripts during foot regeneration at 8 hpa. The plots depict all transcripts that exhibited context-specific expression in untreated samples at 8 hpa. The dotted line demarcates the significance cutoff of FDR ≤ 1e-3. Green dots denote transcripts with significantly diminished injury-induced changes in expression. Orange dots denote transcripts with significantly increased injury-induced changes in expression. (C) Average relative chromatin accessibility plot of peaks containing the TCF TFBM during head and foot regeneration in both untreated and iCRT14-treated samples. iCRT14 treatment significantly diminished and delayed increases in chromatin accessibility associated with the TCF TFBM during head regeneration. (D-F) RNA expression plots showing average log_2_CPM of Wnt signaling transcripts during head and foot regeneration in both untreated and iCRT14-treated samples. iCRT14 treatment prevented or delayed the foot regeneration specific downregulation of injury-induced Wnt signaling genes. (G) Average relative chromatin accessibility plot of peaks containing the AP-1 TFBM during head and foot regeneration in both untreated and iCRT14-treated samples. iCRT14 treatment prolonged the increased chromatin accessibility associated with the AP-1 TFBM (i.e. TPA response element; TRE) during regeneration. (H-J) RNA expression plots showing average log_2_CPM of injury-induced bZIP TFs during head and foot regeneration in untreated and iCRT14-treated samples. iCRT14 treatment prolonged the upregulation of injury responsive bZIP TFs during head and foot regeneration. (K,L) Comparison of the average log_2_FC in (K) transcript abundance or in (L) chromatin accessibility between head and foot regenerates treated with 5 µM iCRT14 at 12 hpa. iCRT14 virtually abolished all context-specific transcription at 8hpa. Features that did not show a significant (FDR ≤ 1e-3) differences in the injury response between head and foot regenerates when compared to 0 hpa controls are highlighted in grey. Features that did show a significant difference are highlighted in red. The dotted line indicates perfect correlation between regeneration types.

Importantly, however, inhibitor treatment did not block the expression of all head regeneration enriched transcripts. Several Wnt pathway IRTs that were part of the context-independent response to injury at 3 hpa in untreated animals—such as *wnt9/10c, wnt3*, and *sp5*—were also upregulated in iCRT14 treated animals (Fig. 3D-F); however, in contrast to untreated animals, these transcripts were not head-specific in iCRT14-treated regenerates at 8 hpa because they were not appropriately downregulated during foot regeneration (Fig. 3B). By 12 hpa, some Wnt signaling-associated transcripts such as *wnt9/10c* and *sp5* began to show signs of asymmetric expression (Fig. 3D,E), while others such as *wnt3* remained upregulated in a context-independent manner (Fig. 3F).

Notably, prolonged expression was not restricted to canonical Wnt signaling genes. We found that several TFs known to have conserved roles in the metazoan wound response were transiently activated during early regeneration in untreated animals but showed prolonged activation when TCF was inhibited. Treatment with iCRT14 led to an increase in the accessibility of the bZIP TF-bound injury-induced TPA response element (TRE) in both head and foot regenerates at 8 and 12 hpa (Fig. 3G). We also found that several injury-responsive bZIP TFs including *creb, jun*, and *fos* were not downregulated appropriately after the initial wound response (Fig. 3H-J). Similarly, the injury responsive Egr TFBM showed an increase in accessibility at 8 and 12 hpa relative to untreated regenerates (supplemental Fig. 2A). These data suggest that aspects of the early wound response are repressed in a TCF-dependent manner during the transition to pre-patterning.

In characterizing the effects of iCRT14 on the expression of individual transcripts, we noticed that seemingly all IRTs that were normally context-specific at 8 hpa were no longer asymmetrically expressed in inhibitor treated animals. We therefore assessed iCRT14’s impact on transcriptional divergence during regeneration genome-wide. At 8 hpa, we found iCRT14 blocked context-specific transcription, with head and foot regenerates appearing nearly indistinguishable (Fig. 3K,L). At 12 hpa, however, we began to see indications of context-dependent transcription in both our chromatin accessibility and transcriptomic data (supplemental Fig. 2B,C). These data therefore demonstrate a global delay in transcriptional specialization and suggest that TCF plays a critical role in the transition from the context-independent wound response to context specific repatterning during head and foot regeneration. This may indicate that the transient activation of Wnt signaling during early foot regeneration is required to potentiate aboral patterning.

### A conserved injury response GRN may directly activate Wnt signaling

Our results suggested that proper regulation of canonical Wnt signaling during regeneration is critical for oral and aboral regeneration. We therefore sought to better understand how the Wnt pathway is regulated during regeneration. The rapid context-independent upregulation of several Wnt pathway components after amputation led us to hypothesize that these genes may be direct targets of the TFs that activate the initial generalized injury response. We investigated this possibility by looking for cis-regulatory elements near Wnt signaling component loci that contained predicting binding sites for injury-responsive TFs. To select TFBMs of interest, we used chromVAR to identify motifs that were associated with significant changes in chromatin accessibility in our ATAC-seq regeneration time course. In addition, we identified candidate genes that were likely to be driving the observed changes in accessibility using two criteria: 1) the presence of a DNA binding domain predicted to bind an injury responsive TFBM, and 2) RNA expression dynamics that correlated with TFBM accessibility.

We identified 32 TFBMs associated with significant variability in chromatin accessibility during the first 12 hours of regeneration (Fig. 4A). A majority of these TFBMs exhibited similar dynamics during both head and foot regeneration, indicating that most early injury responsive TFs were largely unaffected by tissue context. Because Wnt signaling was activated within three hours post-amputation, we subsequently focused on TFBMs that were linked to increases in accessibility from 0 to 3 hpa. We found that the cAMP response element (CRE), which is bound by Jun, Fos, ATF, and CREB bZIP TFs (Eferl and Wagner, 2003), was associated with the largest increase in chromatin accessibility during this interval (Fig. 4B). Consistently, we observed the transient upregulation of *jun, fos*, and two Creb-like IRTs at 3 hpa during both head and foot regeneration (supplemental Fig. 3A-D). These observations suggest that bZIP TFs likely play an important role in triggering the initial response to injury in *Hydra*.

**Figure 4.**
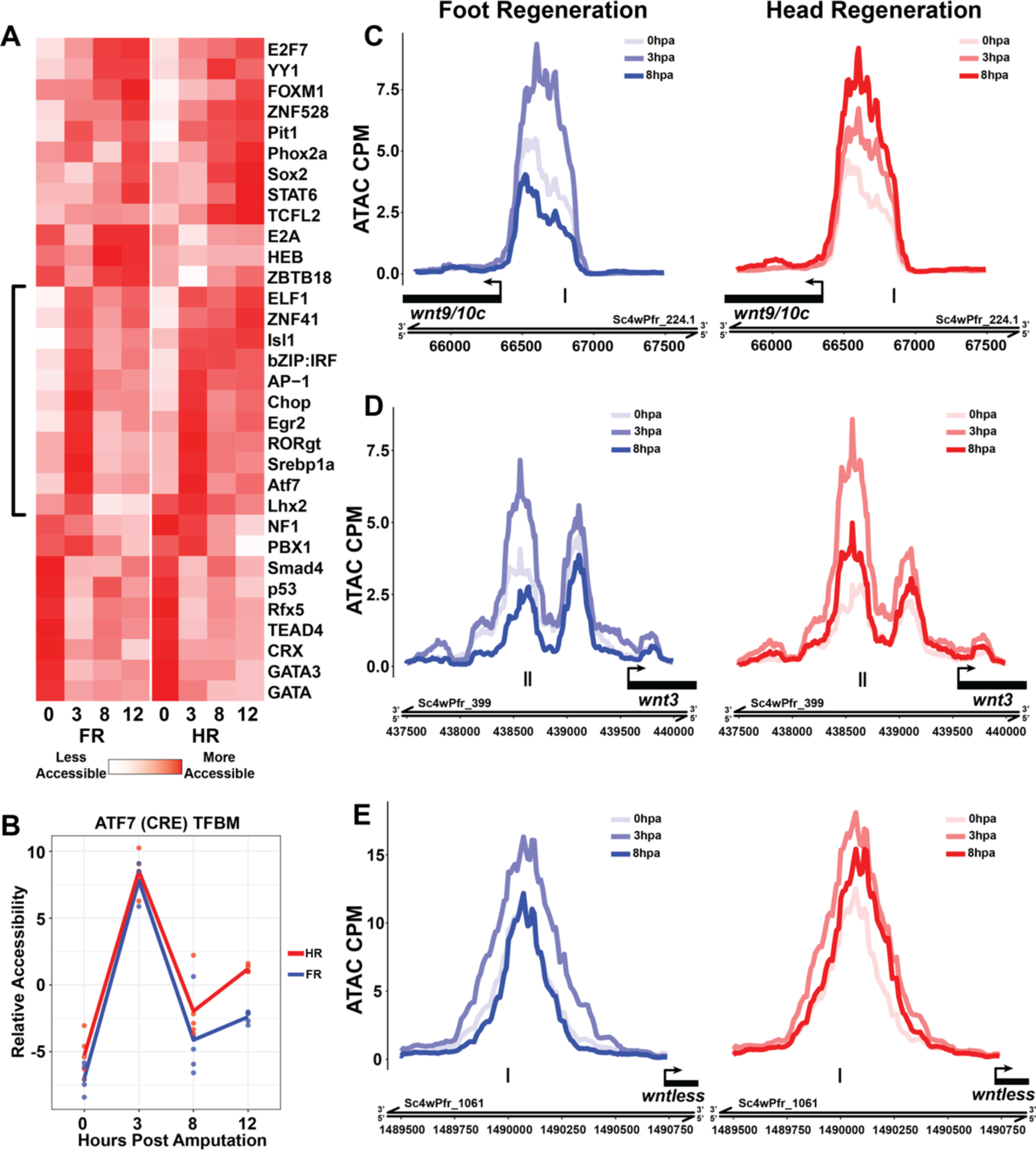
Canonical Wnt signaling genes may be directly activated by injury-responsive transcription factors. (A) Heatmap of the average relative accessibility of TFBMs during regeneration in head and foot regenerates. Transcription factor activity during the first 12 hours of regeneration was highly dynamic and largely similar between head and foot regeneration. TFBMs associated with an increase in accessibility from 0 to 3hpa are demarcated using a square bracket. (B) Average relative chromatin accessibility plot of peaks containing the ATF7 TFBM (i.e. cAMP response elements; CREs) during head and foot regeneration. The ATF7 TFBM is associated with a rapid increase in chromatin accessibility during head and foot regeneration. (C-E) ATAC-seq accessibility data for Wnt signaling gene loci during regeneration. The presumptive promoters of injury-induced wnt signaling genes are likely directly regulated by bZIP transcription factors. Black tick marks indicate predicted CRE hits in putative promoter regions. CPM: average ATAC-seq counts per million calculated using a 10bp bin size.

Next, we analyzed the sequence composition of ATAC-seq peaks near Wnt signaling components to determine if they could plausibly be regulated by injury responsive TFs. We identified multiple predicted instances of the cAMP response element (CRE) in the putative promoters of the injury responsive Wnt pathway genes *wnt9/10c, wnt3*, and *wntless* (Fig. 3C-E). Furthermore, the promoter regions that contained the predicted CRE TFBMs increased in accessibility from 0 to 3 hpa during head and foot regeneration. These data therefore suggest that components of canonical Wnt signaling are direct targets of injury responsive bZip TFs, providing a mechanistic hypothesis for how a patterning pathway is re-activated during regeneration. We propose that this mechanism is the molecular basis for the injury-induced head promoting signal described in the oral induction model.

### Non-amputation injuries activate Wnt signaling and induce ectopic head regeneration if pre-existing organizers are absent

A central principle of the oral induction model is that all injuries produce a transient pulse of head promoting activity, and that the lack of new head organizer formation after certain injuries is due to extrinsic signals from the surrounding tissue context. We therefore reasoned that we could cause a non-amputation injury to undergo head regeneration by experimentally shifting the balance between injury-induced Wnt signaling and the inhibitory signals produced by the surrounding tissue.

To stimulate high levels of injury-induced Wnt signaling, we created a constitutively active injury signal by transversely impaling *Hydra* on fishing line at the midpoint of their oral-aboral axis (Fig. 5A). This created two large puncture wounds that could not close while the fishing line remained in place. To assess the effects of prolonging the injury response, we either removed the fishing line immediately after impalement (0-hour timepoint) or left it in place for 12 hours. To create a permissive and positionally unbiased tissue context, we removed both the head and foot organizers just prior to impalement. We then assessed wound repair outcomes four days post-injury.

**Figure 5.**
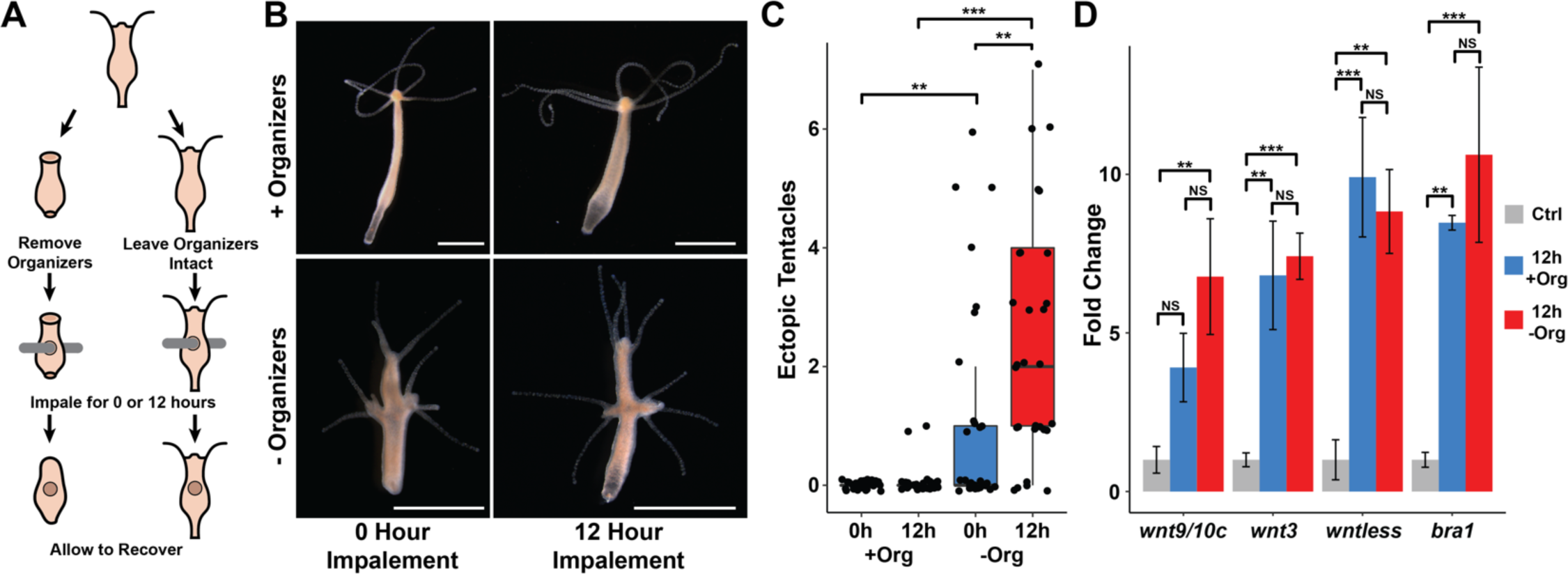
Non-amputation injuries induce ectopic head regeneration when pre-existing organizers are removed. (A) Experimental design to test for the sufficiency of injury to induce new organizer formation in tissue with or without pre-existing organizers. (B) Prolonged injuries induce ectopic head regeneration when pre-existing organizers are removed. Animals were imaged 4 days post-injury, giving sufficient time for removed head and foot structures to regenerate. (C) Quantification of the number of ectopic tentacles induced by various impalement injury conditions. Both the presence of pre-existing organizers and the duration of the injury signal significantly contributed to the ectopic head regeneration phenotype. -Org indicates that organizers were removed just prior to impalement, +Org indicates organizers were left intact. (D) RT-qPCR results for head-specific transcripts in tissue that had been impaled for 12 hours. The Wnt signaling genes *wnt3, wntless*, and *wnt9/10c* and the head-specific transcription factor *bra1* are significantly upregulated after 12 hours of impalement. Uninjured body column tissue was used as a control. Significance was evaluated using ANOVA and Tukey’s HSD test. Error bars indicate standard deviation. All statistical analyses were performed on 2^-ΔCq^ values. HR: head regeneration, FR: foot regeneration, * indicates P-value ≤ 0.05, ** indicates P-value ≤ 0.01, *** indicates P-value < 0.001.

In polyps with intact organizers, impalement never resulted in secondary axes. However, when organizers were removed prior to injury, impalement resulted in gross patterning defects. Ectopic head structures—ranging from single tentacles to fully formed heads—formed at one or both of the presumptive impalement sites in ∼83% of cases (24/29; Fig. 5B). The extent of this phenotype was dependent on the duration of the injury signal, as animals that were impaled and then immediately allowed to recover grew significantly fewer ectopic tentacles than animals that had been impaled for 12 hours (Fig. 5C). We noted that this phenotype was strain-dependent: we observed ectopic head regeneration after impalement when using Basel and 105 strain animals, but not when using AEP strain animals (Supplemental Fig. 4A-C). We speculate that this may be caused by genetic variation affecting inhibition levels in body column tissue.

Using qPCR, we found that the Wnt signaling genes *wnt9/10c, wnt3*, and *wntless* as well as the head specific TF *brachyury1* were significantly upregulated after 12 hours of impalement (Fig. 5D). Notably, the upregulation of *wntless, wnt3*, and *brachyury1* was context-independent, as their expression at 12 hours post impalement was not affected by the presence of pre-existing organizers. This is in contrast to head and foot regeneration following mid-gastric bisection, where the expression of these transcripts was context-specific by 12 hpa. Thus, preventing wound closure prolonged the injury-induced context-independent phase of expression for several head specific transcripts. In contrast, *wnt9/10c* appeared to show increased upregulation in the absence of pre-existing organizers as compared to when organizers were present, however this differential fell just outside our significance threshold (P = 0.06).

These data, in conjunction with our analysis of transcription during head and foot regeneration, strongly support the oral induction model and demonstrate that Wnt signaling is activated in wounded body column tissue irrespective of injury type (i.e. amputation or non-amputation) or context (i.e. with or without pre-existing organizers). Furthermore, these findings show that injuries that are typically non-regenerative are capable of inducing the development of a fully functional head organizer, but only do so when the surrounding tissue context is permissive.

## Discussion

Based on our comprehensive characterization of transcription during the first 12 hours of head and foot regeneration, we propose the following model of patterning during regeneration in *Hydra* (Fig. 6): injury-induced bZIP TFs activate a head specifying program through the upregulation of Wnt ligands, regardless of the injury’s tissue context. This triggers a Wnt-based feed-forward loop that promotes the formation of a new head organizer. However, if a head organizer is already present, Wnt signaling is inhibited and a foot-specific program is activated instead. This model is highly consistent the oral induction model, and thus provides a molecular basis for the theoretical framework established by earlier research.

**Figure 6.**
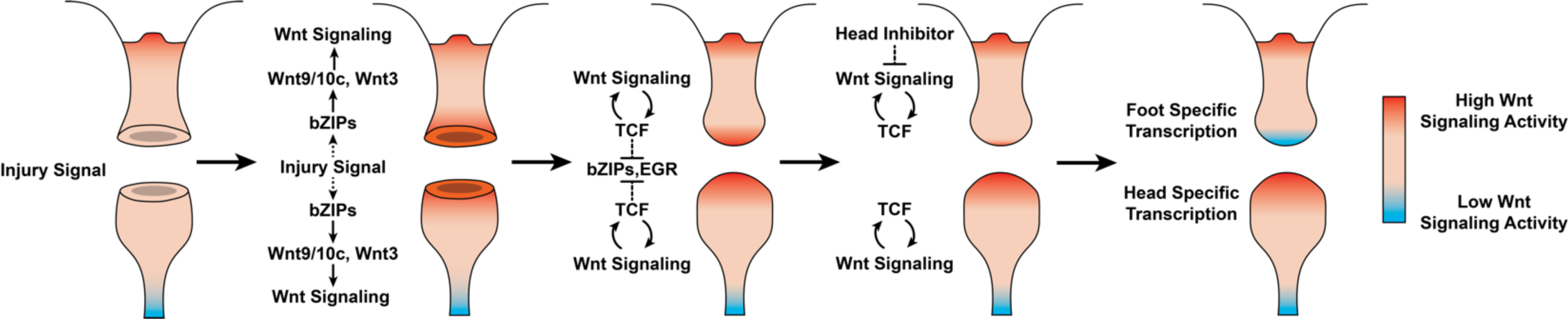
Proposed model of positional decision making following mid-gastric bisection in *Hydra*. Injury triggers a Wnt signaling cascade regardless of tissue context through the direct upregulation of the Wnt ligands *wnt9/10c* and *wnt3* via injury-responsive bZIP transcription factors. An increase in Wnt signaling then represses injury-responsive transcription factors in a TCF-dependent manner. While Wnt signaling has the capacity for autocatalytic amplification, this only occurs during head regeneration, as the autocatalytic amplification is blocked during foot regeneration by an uncharacterized inhibitor originating from pre-existing oral tissue. After Wnt signaling has been repressed in foot regenerates, foot specific transcription is activated. HR: head regeneration, FR: foot regeneration. Red coloration denotes high Wnt signaling activity. Blue coloration denotes low Wnt signaling activity.

Similar to what has been reported in other highly regenerative animals (DuBuc et al., 2014; Schaffer et al., 2016; Wenemoser et al., 2012; Wurtzel et al., 2015), we found that the initial transcriptional response to injury in *Hydra* was the same regardless of the surrounding tissue context. This generalized wound response included the activation of the canonical Wnt pathway, which provides a molecular basis for the transient increase in the capacity of injured tissue to form a new head organizer (Kobatake and Sugiyama, 1989; MacWilliams, 1983a; Müller, 1996). Our data suggests that this increase in Wnt signaling is driven by injury-responsive bZIP TFs that directly upregulate Wnt pathway components, including *wnt9/10c, wnt3*, and *wntless*. Notably, a similar hypothesis had been proposed previously based on the presence of predicted bZIP binding sites in the *wnt3* promoter and the observation that Creb is transiently activated during early regeneration in *Hydra* (Galliot et al., 1995; Nakamura et al., 2011). Our chromatin accessibility data strongly support the importance of bZIP TFs in the initial response to injury in *Hydra* and suggest that additional Wnt signaling components beyond *wnt3* are directly targeted by bZIP TFs.

Further supporting this hypothesis, Tursch and colleagues report that mitogen activated protein kinases (MAPKs)—direct upstream regulators of bZIP TFs—are rapidly activated by injury in a context and position-independent manner via Ca+ and ROS signaling (Tursch et al., 2020). Disrupting this injury-induced activation of MAPKs resulted in a significant downregulation of *wnt9/10c* and blocked both head and foot regeneration. However, although inhibition of the MAPK Erk1/2 resulted in a significant downregulation of *wnt3* during regeneration, inhibition of p38 and JNK caused a significant increase in *wnt3* expression. These findings provide strong evidence implicating bZIP TFs as regulators of the canonical Wnt signaling pathway during the wound response and suggest that *wnt3* and *wnt9/10c* may have distinct roles during regeneration and may each be regulated by a distinct set of injury-responsive TFs. The complexity underlying the injury-responsive activation of Wnt signaling highlights the need for future research to functionally characterize the roles of Wnt9/10c and Wnt3 during regeneration and to identify the direct targets of injury-responsive bZIP TFs.

We found that injury-induced Wnt signaling became restricted to head regeneration between 3 and 8 hpa. We hypothesize that this divergence is driven by the balance of two competing processes: the autoregulatory amplification of injury induced Wnt signaling and the long-range repression of Wnt signaling by head tissue. This is supported by our finding that non-amputation injuries induced Wnt signaling and could trigger ectopic head formation when the source of inhibition (i.e. head tissue) was removed. Currently, the molecular basis for this non-cell autonomous inhibition is poorly understood. Sp5 was recently identified as a head inhibitor in *Hydra* (Vogg et al., 2019), but, as it is a transcription factor, it is unlikely to act in a non-cell autonomous manner. However, we found that the secreted Wnt inhibitor *notum* was enriched in uninjured head tissue in the *Hydra* scRNA-seq atlas (supplemental Fig. 5A) (Siebert et al., 2019). We therefore hypothesize that Wnt signaling is inhibited during foot regeneration by extracellular *notum* emanating from the pre-existing head organizer.

Although the oral induction model provides a well-defined mechanism for head specification during regeneration, the process of foot specification remains far less clear. Mature foot tissue possesses organizing capacity and produces inhibitory signals that block ectopic foot formation (Cohen and MacWilliams, 1975; J Hicklin and Wolpert, 1973; MacWilliams and Kafatos, 1968), but none of the molecular pathways underlying these phenomena have been identified. Unlike head regeneration, we did not find evidence of injury-induced context-independent signaling pathways or regulatory genes that became restricted to foot regeneration at later timepoints. Rather, we found that foot specific transcription was activated after the generic increase in Wnt signaling had stopped. Thus, the inhibition of injury-induced Wnt signaling by pre-existing head tissue may be an important early step in potentiating foot specification during regeneration. This is consistent with the observation that foot regeneration is slower in headless tissue fragments, as regenerating heads produce less inhibition than mature head tissue (Grens et al., 1996; MacWilliams, 1983b; Müller, 1995). Notably, TCF inhibition blocked the activation of both head and foot specific transcription during regeneration, suggesting that TCF is required for the re-specification of both axial identities. This may be linked to the failure to downregulate the generalized wound response when TCF is inhibited. However, because the direct targets of TCF are largely uncharacterized in *Hydra* and because we cannot eliminate the possibility of off-target effects from iCRT14 treatment, additional research is needed to understand the role of TCF during foot regeneration.

Regenerative capacity is broadly but irregularly distributed across metazoan phyla (Brockes et al., 2001; Sanchez Alvarado and Tsonis, 2006). Therefore, identifying both the conserved and derived aspects of wound repair across diverse model systems can provide insight into the evolution of the injury response in animals. Activation of canonical Wnt signaling appears to be a near universal hallmark of the injury response in animals and is required for wound repair in both regenerative and non-regenerative species (Whyte et al., 2012). Such a deeply conserved role suggests that regulatory mechanisms connecting Wnt signaling to the wound response predate the split of cnidarians and bilaterians over 500 million years ago. This raises the possibility that the injury responsive GRNs that activate Wnt signaling may also be conserved throughout the animal kingdom. A conserved role for bZIPs in activating Wnt signaling during regeneration is supported by the findings that Erk is required for the injury-induced activation of Wnt signaling in planarians (Owlarn et al., 2017) and bZIP TFBMs are required for Wnt expression during imaginal wing disc regeneration in *Drosophila* (Harris et al., 2016). In contrast, it was recently reported that the zinc finger TF Egr is required for Wnt expression during acoel regeneration (Ramirez et al., 2020). It will therefore be important for future research to characterize injury-responsive GRNs across diverse animal taxa to better understand the evolutionary history of Wnt signaling and its regulation during wound repair.

## Materials and Methods

### Code and Data Availability

All code used in this study is available both as a git repository at github.com/cejuliano/jcazet_regeneration_patterning and on Dryad at doi.org/10.25338/B8S612. FASTQ files of raw ATAC-seq and RNA-seq reads, expression matrices for ATAC-seq and RNA-seq reads mapped to the *Hydra* 2.0 genome reference, consensus peak files, and bigwig genome tracks of individual and pooled ATAC-seq replicates are available through the Gene Expression Omnibus under the accession GSE152994. The Hydra 2.0 genome gene model IDs associated with the gene names used throughout this study are provided in Supplemental Table 3.

### ATAC-seq Library Preparation

For each ATAC-seq replicate, ∼15 whole, bud-free *Hydra vulgaris* (strain 105) polyps that had been fed once weekly were starved for two days and then transversely bisected at the midpoint of their oral-aboral axis. Regeneration was then allowed to proceed for 0, 3, 8, or 12 hours. Regenerating tips corresponding to ∼1/3 of the total regenerate length were then isolated from head and foot regenerates and used for generating ATAC-seq libraries. For iCRT14 treatment, 10 mM iCRT14 (Sigma-Aldrich, St. Louis, MO SML0203) dissolved in DMSO was diluted in *Hydra* medium to a final concentration of 5 µM. iCRT14 solution was added to the animals 2 hours before amputation and was left in place until the tissue was collected for library preparation.

To generate ATAC-seq libraries, we made use of a modified version of the OMNI-ATAC protocol as described previously (Corces et al., 2017; Siebert et al., 2019). Briefly, regenerating tips were washed once with 1 ml of chilled *Hydra* dissociation medium (DM) (Gierer et al., 1972) and then homogenized in 1ml fresh DM using ∼30 strokes of a tight-fitting dounce. Cells were then pelleted at 500 RCF for 5 minutes at 4°C in a benchtop centrifuge and subsequently lysed in 50ul of chilled resuspension buffer (RSB; 10 mM Tris-HCl -pH 7.4, 10 mM NaCl, 3 mM MgCl2) with 0.1% Tween-20, 0.1% NP-40, and 0.01% digitonin for 3 minutes on ice. Lysis was halted by adding 1 ml of RSB with 0.1% Tween-20. The lysate’s nuclear concentration was then quantified by loading 19 ul of lysate and 1 ul of 0.2 mg/ml Hoechst 33342 onto a Fuchs-Rosenthal hemocytometer. A volume of lysate corresponding to ∼50,000 nuclei was then aliquoted and spun at 500 RCF for 10 minutes at 4°C in a benchtop centrifuge. The resulting crude nuclear pellet was then resuspended in 50 ul of tagmentation solution (1x TD buffer (Illumina, San Diego, CA 20034197), 33% PBS, 0.01% digitonin, 0.1% tween-20, 5 ul TDE1 (Illumina 20034197)) and shaken at 1000 RPM for 30 minutes at 37°C. Tagmentation was halted by adding 250 ul of PB buffer from a Qiagen MinElute PCR Purification Kit (Qiagen, Hilden, Germany 28004).

Tagmented DNA was purified using a Qiagen MinElute PCR Purification Kit using the standard manufacturer’s protocol. Libraries were then amplified with 2X NEBNext master mix (NEB, Ipswitch, MA M0541S) using cycle numbers determined by qPCR as described in the standard ATAC-seq protocol (Buenrostro et al., 2015, 2013). Agencourt AMPure XP beads (Beckman Coulter, Pasadena, CA A63881) were then used to purify libraries and restrict fragment sizes to between 100 and 700 base pairs. Libraries were then pooled and sequenced on an Illumina HiSeq4000 using 2×150bp reads.

### ATAC-seq Data Processing

Sequencing adapters, stretches of low-quality base calls, and unpaired reads were removed from the raw sequencing data using Trimmomatic (Bolger et al., 2014). Filtered reads were then mapped to the *Hydra vulgaris* 2.0 genome (arusha.nhgri.nih.gov/hydra/) using Bowtie2 (Langmead and Salzberg, 2012). Mitochondrial reads were removed by independently mapping filtered reads to the *Hydra* mitochondrial genome (Voigt et al., 2008) and removing mitochondrial reads from the genome-mapped data using Picard Tools (broadinstitute.github.io/picard). Ambiguously mapped reads (defined as having a MAPQ value of ≥ 3) and discordantly mapped read pairs were removed using SAMtools (Li et al., 2009). PCR duplicates were labeled using Picard Tools and removed using SAMtools.

Peak calling was performed using a modified version of the ENCODE consortium’s ATAC-seq analysis pipeline (encodeproject.org/atac-seq) (Landt et al., 2012). First, unambiguously mapped deduplicated non-mitochondrial reads were centered over the transposon binding site by shifting + strand reads +4bp and – strand reads -5bp using deepTools2 (Ramírez et al., 2016). Peaks were then called using Macs2 (Zhang et al., 2008) with a permissive p-value cutoff of 0.1. We then generated consensus lists of biologically reproducible peaks using the irreproducible discovery rate (IDR) framework (Li et al., 2011) by identifying peaks that were reproducible (IDR score ≥ 0.1) across at least three pairwise comparisons of biological replicates in at least one treatment group.

To assess the quality and reproducibility of our ATAC-seq data, we made use of several metrics used by the ENCODE consortium. Because core promoters are expected to be highly accessible, ATAC-seq reads should be strongly enriched near transcription start sites (TSS). The current *Hydra* genome gene models lack UTR annotations, so we determined the TSS enrichment score for each biological replicate by measuring read enrichment near the first start codon of the 2000 most highly expressed genes in the *Hydra* single cell RNA-seq atlas relative to the average read density ± 1 kb from the start codon. We observed between a 5- and 8-fold TSS enrichment across the individual samples in our dataset. We also calculated the self-consistency and rescue ratios for each treatment group to evaluate consistency across replicates. We found that all treatment groups had self-consistency and rescue ratios less than 2, indicating good overall reproducibility across replicates. A full table of ATAC-seq library metrics are provided in Supplementary Table 1.

### Differential Accessibility and ChromVar Analysis

Read counts for peaks in our consensus peakset were calculated using the R Diffbind package (Ross-Innes et al., 2012). EdgeR was then used for downstream differential accessibility analyses (Robinson et al., 2010). Peaks that did not have at least 10 counts per million in at least 3 replicates were excluded from further analyses. Differentially accessible peaks were then identified using a quasi-likelihood negative binomial generalized log-linear model with a False Discovery Rate (FDR) cutoff of 1e-4. The full results tables for all ATAC-seq pairwise comparisons performed as part of the edgeR analysis are included in Supplementary File 1.

For the chromVAR TFBM accessibility analyses, the width of all peaks that showed a significant change during regeneration was fixed at 250bp and new read counts were generated. Changes in accessibility were then calculated for all TFBMs provided in the custom list of HOMER motifs found in the chromVARmotifs package. Because our unfiltered enrichment results often included redundant TFBMs with highly similar sequence composition and accessibility dynamics, TFBM redundancy was reduced by performing hierarchical clustering on a matrix of pairwise TFBM similarity scores generated by the HOMER compareMotifs function. The TFBM within a cluster that showed the greatest change in accessibility across treatments was then used as the representative motif for that cluster.

### RNA-seq Library Preparation

30 whole, bud-free *Hydra vulgaris* (strain 105) polyps that had been fed once weekly were starved for two days and then transversely bisected at the midpoint of their oral-aboral axis in batches of 10 to reduce variability in amputation and collection times. Regeneration was then allowed to proceed for 0, 3, 8, or 12 hours. Regenerating tips corresponding to ∼1/3 of the total regenerate length were then isolated from head and foot regenerates and frozen in Trizol (Thermo Fisher Scientific, Waltham, MA 15596018) at -80°C. The three individual batches of 10 regenerating tips were then pooled into a single biological replicate and RNA was subsequently extracted using a standard Trizol RNA purification protocol. DNA contamination was removed using the Qiagen DNase Set (79254) following the manufacturer’s protocol. A final cleanup was then performed using a Zymogen RNA Clean and Concentrator kit (Zymogen, Irvine, CA R1017) following the standard manufacturer’s protocol. Strand-specific polyA-enriched libraries were prepared using the Kapa mRNA-seq Hyper kit (Kapa Biosystems, Cape Town, South Africa KK8581). Untreated samples were sequenced on an Illumina HiSeq4000 with 1×100bp reads. 12 hpa and all iCRT14 treated samples were sequenced on an Illumina NovaSeq using 2×150bp reads.

### RNA-seq Data Processing and Differential Gene Expression Analysis

Sequencing adapters and stretches of low-quality base calls were removed from the raw sequencing data using Trimmomatic. Filtered reads were then mapped to the *Hydra* 2.0 gene models and read counts per gene were calculated using RSEM (Li and Dewey, 2011). A full table of RNA-seq library metrics are provided in Supplementary Table 1. Using edgeR, read counts were normalized and genes that did not have at least 2 counts per million in at least 3 replicates were excluded from further analyses. Differentially expressed genes were then identified using a quasi-likelihood negative binomial generalized log-linear model with a False Discovery Rate (FDR) cutoff of 1e-3. The full results tables for all RNA-seq pairwise comparisons performed as part of the edgeR analysis are included in Supplementary File 2.

### Hydra Tissue Manipulation Experiments

*Hydra vulgaris* (either Basel, 105, or AEP strain) polyps that had been fed twice weekly were starved for at least 24 hours and then impaled at the midpoint of their oral-aboral axis in batches of ∼15 on a single piece of 0.3 mm diameter fishing line. To remove pre-existing organizer tissue before impalement, heads and feet were removed by performing transverse amputations just below the tentacle ring and just above the peduncle respectively. The *Hydra* were then either removed from the fishing line immediately after being impaled for the 0-hour timepoint or were left on the fishing line for 12 hours and then removed. Phenotypes were then documented after four days of recovery post-impalement. Significant differences in the number of ectopic tentacles was determined using an ANOVA followed by Tukey’s HSD test in R using the agricolae package.

### Quantitative Reverse Transcription PCR (qPCR)

Impaled *Hydra* were prepared as described above, with 15 polyps per biological replicate and a total of 3 biological replicates per treatment. Tissue surrounding the impalement site, corresponding to ∼1/4 of the total polyp length, was isolated via amputation and frozen in Trizol at -80°C. RNA was isolated using a standard Trizol extraction. Contaminating DNA was removed using the Qiagen DNase Set following the manufacturer’s protocol. DNaseI and residual contamination was then removed using a Zymogen RNA Clean and Concentrator kit following the manufacturer’s protocol. cDNA was synthesized using 1 µg of purified RNA and Promega M-MLV RNase H Minus Point Mutant Reverse Transcriptase (Promega, Madison, WI M3682) using the manufacturer’s recommended protocol for oligo dT-primed synthesis. cDNA was then diluted 1:3 in nuclease free water for use in qPCR experiments.

Three technical replicates of 10 ul qPCR reactions per sample were prepared using Bio-Rad SsoAdvanced universal SYBR green master mix (Bio-Rad, Hercules, CA 1725271) and were run on a CFX96 Touch Real-Time PCR Detection System (Bio-Rad 1855195). Cq values from technical replicates were pooled for subsequent analyses. *rp49* was used as an internal control to calculate ΔCq values after first being found to give similar results across all treatments when compared to a second housekeeping gene, *actin*. Statistically significant differences in mRNA expression was calculated by performing an ANOVA analysis on 2^-ΔCq^ values followed by Tukey’s HSD test in R using the agricolae package.

## Supporting information

Supplementary_File_1-ATAC_DGE_Tables

Supplementary_File_2-RNA_DGE_Tables

## Acknowledgements

We thank Bruce Draper, Gary Wessel, Rob Steele, Bryan Teefy, Sergio Campos, Ben Cox, Thomas Holstein, Anja Tursch, Prashanth Rangan, Jeffrey Farrell, and Stefan Materna for critical reading of the manuscript; the members of the Juliano lab for their thoughtful input on this study; Vanessa Rashbrook, Emily Kumimoto, Siranoosh Ashtari, and Lutz Froenicke from the DNA Technologies and Expression Analysis Core at the UC Davis Genome Center (supported by NIH Shared Instrumentation Grant 1S10OD010786-01) for technical advice and assistance with library preparation and sequencing; and the Vincent J. Coates Genomics Sequencing Laboratory at UC Berkeley for additional library preparation and sequencing. Funding support for this study was provided by UC Davis start-up funds and the NIH grant R35 GM133689 awarded to CEJ.

