## Supplementary Material for "Oral Regeneration Is the Default Pathway Triggered by Injury in *Hydra*"

### Supplemental Figures

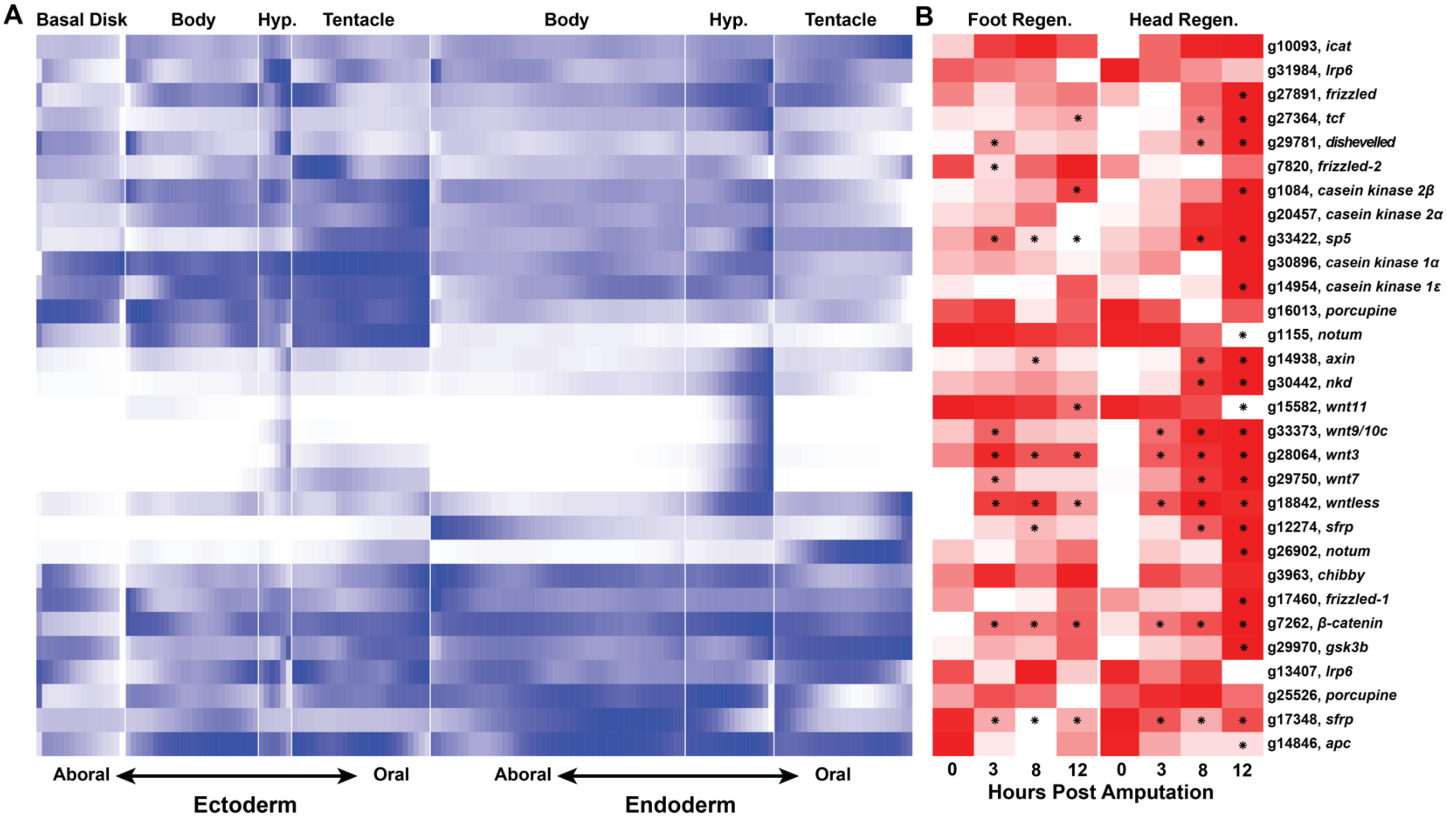


**Supplemental Figure 1. Several head specific canonical Wnt signaling components are upregulated during head and foot regeneration.** (A) Heatmap depicting canonical Wnt signaling component expression in positionally ordered single cell RNA-seq data. Wnt signaling components exhibit a diverse set of expression patterns in epithelial cell lineages. The expression of Wnt signaling components along the oral-aboral axis was visualized using the results of a previously published analysis (Siebert et al., 2019) that predicted the positional order of epithelial cells along the oral-aboral axis in the *Hydra* single cell atlas. (B) Heatmap of canonical Wnt signaling components during regeneration. Several canonical Wnt signaling genes were transcriptionally upregulated during early foot regeneration. Darker coloration indicates higher expression. Asterisks indicate a significant change (FDR ≤ 1e-3) in RNA abundance relative to 0 hpa controls. Hyp: hypostome.

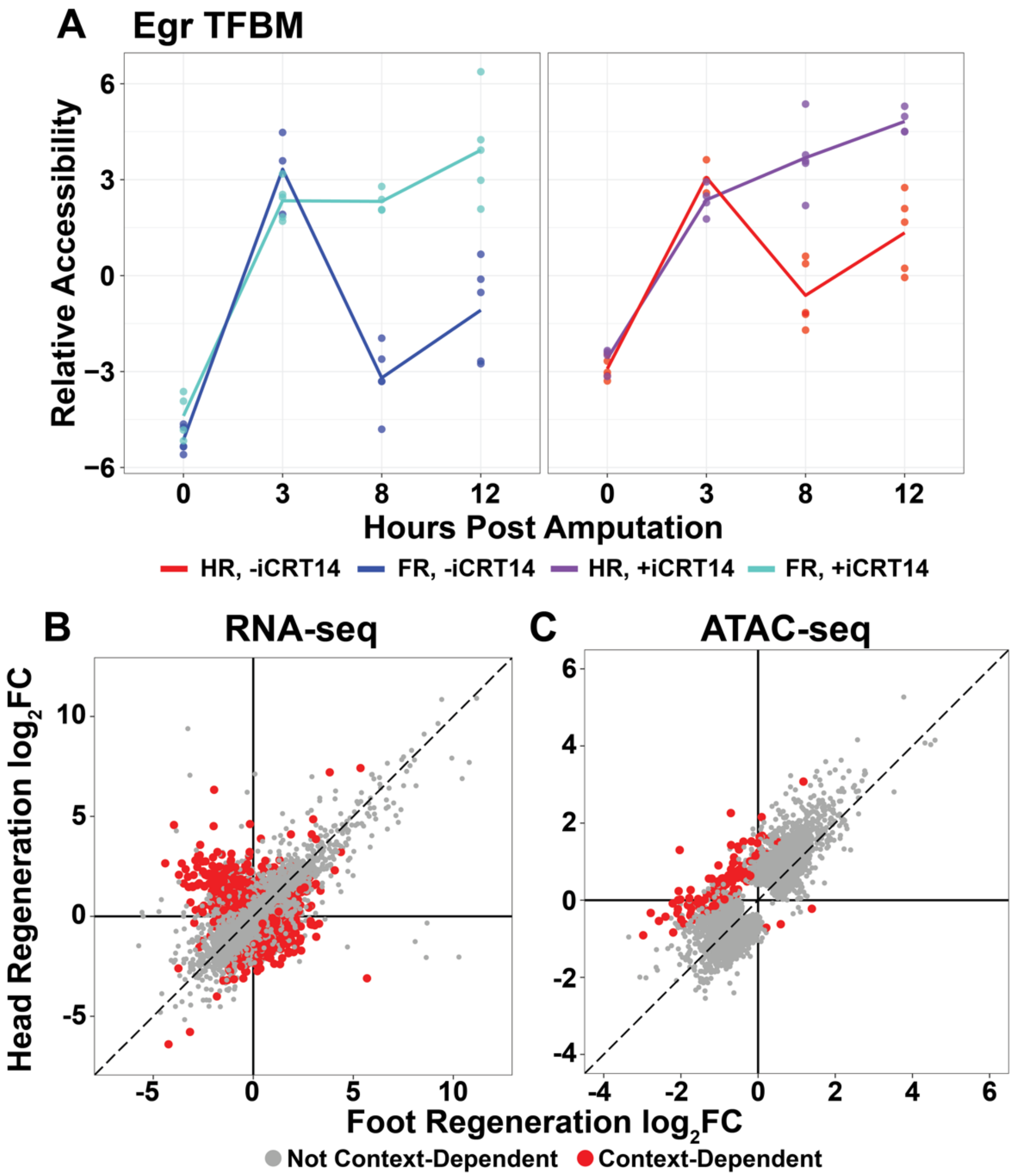


**Supplemental Figure 2. iCRT14 delays the transition from the context-independent wound response to pre-patterning.** (A) Average relative chromatin accessibility plot of peaks containing the Egr TFBM during head and foot regeneration. The injury induced increase in Egr TFBM accessibility was prolonged in the presence of iCRT14. (B,C) Comparison of the average $log_2FC$ in (B) transcript abundance or in (C) chromatin accessibility between head and foot regenerates treated with 5 μM iCRT14 at 12 hpa. iCRT14-treated regenerating tissue showed evidence of context-dependent transcription by 12 hpa. Features that did not show a significant (FDR ≤ 1e-3) differences in the injury response between head and foot regenerates when compared to 0 hpa controls are highlighted in grey. Features that did show a significant difference are highlighted in red. The dotted line indicates perfect correlation between regeneration types.

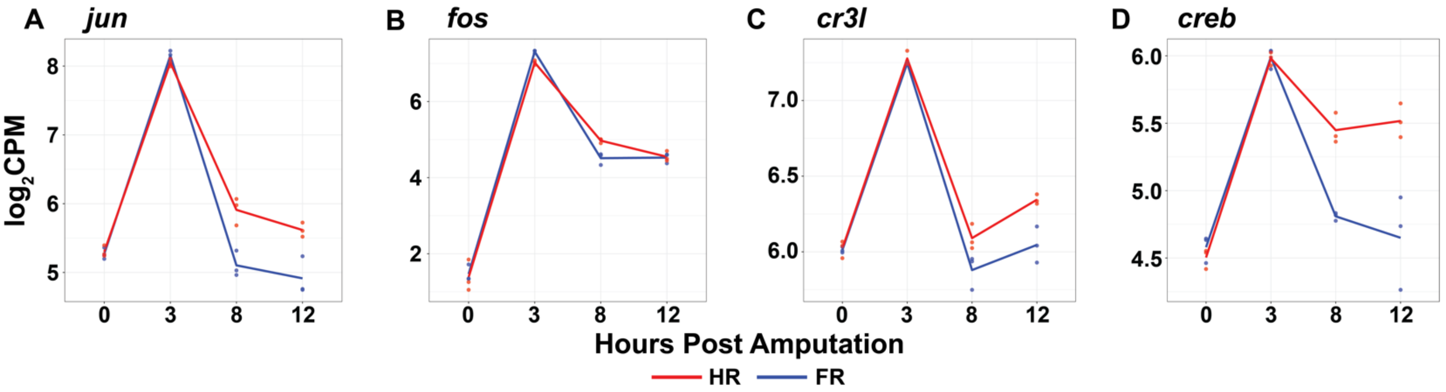


**Supplemental Figure 3. Basic leucine zipper transcription factors are transiently upregulated during head and foot regeneration at 3 hpa.** (A-D) RNA expression plots showing average normalized RNA-seq read counts for bZIP TFs in $\log_2$ counts per million ($\log_2$CPM) during head and foot regeneration.

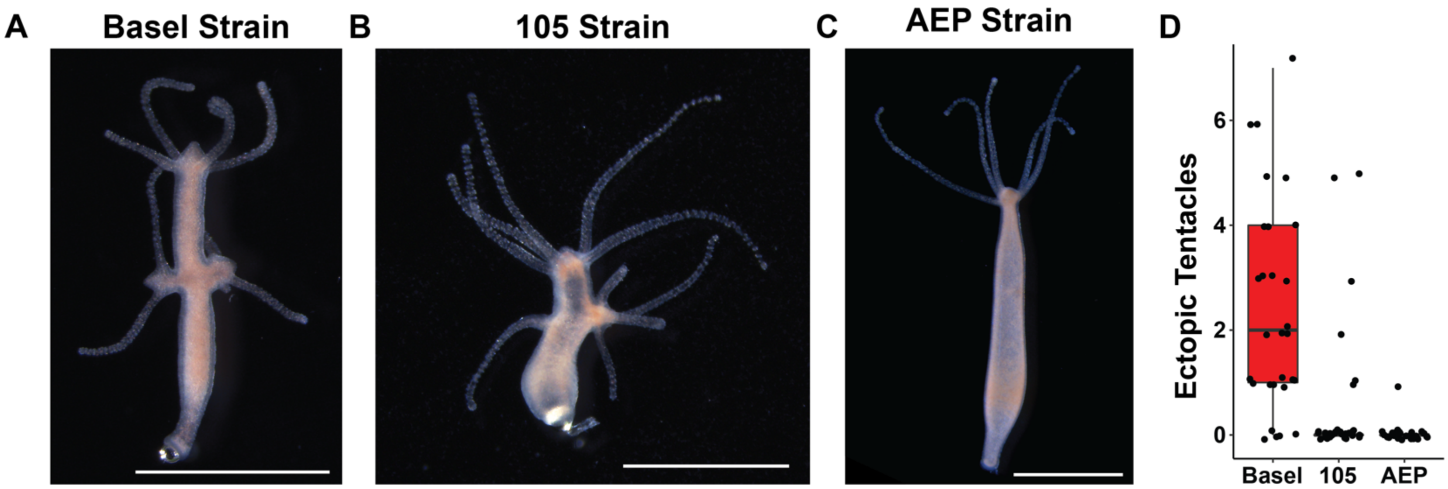


**Supplemental Figure 4. Ectopic head regeneration induced by impalement injuries is strain dependent.** (A-C). *Hydra* polyps lacking pre-existing organizers were impaled for 12 hours on fishing line then allowed to recover for 4 days. Basel (A) and 105 (B) strain *Hydra* exhibit ectopic head regeneration following impalement, but AEP strain *Hydra* (C) do not. (D) Quantification of the number of ectopic tentacles induced by 12 hours of impalement for each strain.

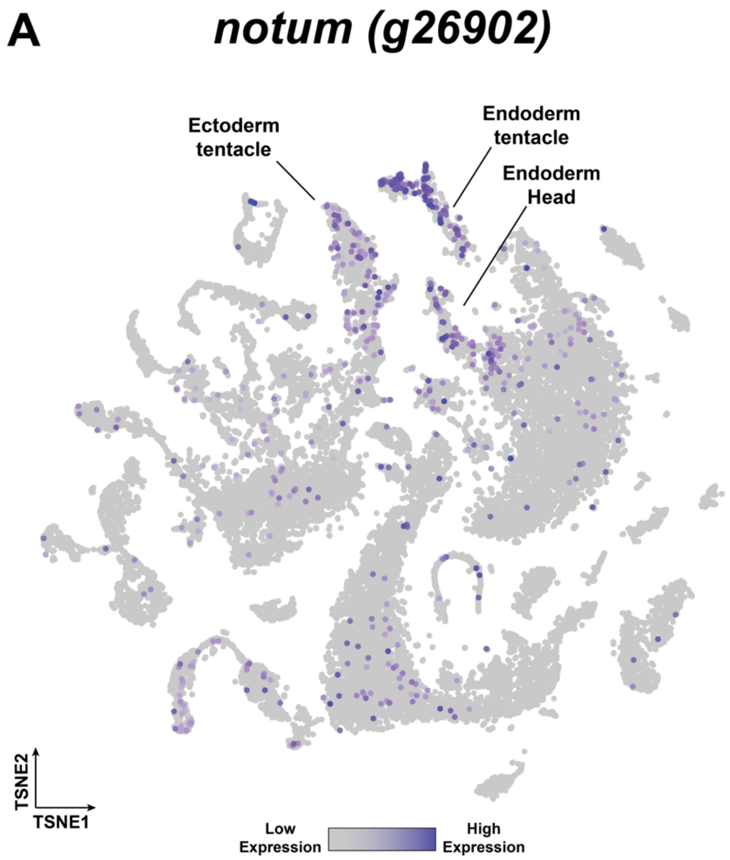


**Supplemental Figure 5. *notum* is expressed in head tissue in whole uninjured *Hydra*.** (A) t-distributed Stochastic Neighbor Embedding (t-SNE) plot depicting notum expression in the *Hydra* single cell RNA-seq atlas.

#### Supplementary Table 1

| Sample ID | HPA | iCRT | Total Read Pairs | Final Mapped Read Pairs | TSS Enrichment | Reproducible Peaks | Self-Consistency Ratio | Rescue Ratio |
|---|---|---|---|---|---|---|---|---|
| 0F1 | 0 | - | 68865654 | 31845047 | 6.15 | 78158 | 1.17 | 1.00 |
| 0F2 | 0 | - | 65301503 | 21970963 | 7.16 | | | |
| 0F3 | 0 | - | 80642153 | 25843928 | 6.61 | | | |
| 0F4 | 0 | - | 76300368 | 25925595 | 6.79 | | | |
| 0F5 | 0 | - | 81363105 | 20714765 | 6.79 | | | |
| 0H1 | 0 | - | 72424905 | 31676679 | 6.31 | 82396 | 1.17 | 1.04 |
| 0H2 | 0 | - | 68870186 | 27376967 | 6.81 | | | |
| 0H3 | 0 | - | 81121398 | 31084966 | 6.50 | | | |
| 0H4 | 0 | - | 67378757 | 26076124 | 6.58 | | | |
| 0H5 | 0 | - | 81380944 | 26247390 | 6.44 | | | |
| 3F1 | 3 | - | 120749030 | 35747201 | 6.44 | 57980 | 1.11 | 1.20 |
| 3F2 | 3 | - | 133453367 | 37551124 | 6.85 | | | |
| 3F3 | 3 | - | 128451210 | 44807082 | 6.76 | | | |
| 3H1 | 3 | - | 127697432 | 33201124 | 7.45 | 58259 | 1.09 | 1.15 |
| 3H2 | 3 | - | 124784335 | 38517160 | 7.09 | | | |
| 3H3 | 3 | - | 130420684 | 36703497 | 7.05 | | | |
| 8F1 | 8 | - | 71642160 | 20270917 | 7.30 | 84248 | 1.29 | 1.05 |
| 8F2 | 8 | - | 77325666 | 17968245 | 7.83 | | | |
| 8F3 | 8 | - | 75894343 | 17391339 | 7.15 | | | |
| 8F4 | 8 | - | 71117490 | 25653837 | 6.98 | | | |
| 8F5 | 8 | - | 74203082 | 30614424 | 7.17 | | | |
| 8H1 | 8 | - | 72411934 | 18714649 | 7.18 | 83974 | 1.18 | 1.04 |
| 8H2 | 8 | - | 76210411 | 16618719 | 7.78 | | | |
| 8H3 | 8 | - | 74495918 | 22207972 | 6.53 | | | |
| 8H4 | 8 | - | 68874107 | 28397085 | 6.78 | | | |
| 8H5 | 8 | - | 85121623 | 26333487 | 6.31 | | | |
| 12F1 | 12 | - | 35989738 | 16371364 | 7.81 | 71949 | 1.09 | 1.00 |
| 12F2 | 12 | - | 37815306 | 16333732 | 7.26 | | | |
| 12F3 | 12 | - | 31696505 | 10175518 | 7.47 | | | |
| 12F4 | 12 | - | 35835498 | 10383772 | 8.43 | | | |
| 12F5 | 12 | - | 27753392 | 9458838 | 7.78 | | | |
| 12H1 | 12 | - | 46530322 | 17225290 | 6.98 | 71463 | 1.16 | 1.05 |
| 12H2 | 12 | - | 33411133 | 16231389 | 6.68 | | | |
| 12H3 | 12 | - | 38128076 | 9053148 | 8.13 | | | |
| 12H4 | 12 | - | 37038785 | 8069733 | 7.94 | | | |
| 12H5 | 12 | - | 35508148 | 9077295 | 7.10 | | | |
| 0iF1 | 0 | + | 29450456 | 13933591 | 6.99 | 68393 | 1.13 | 1.03 |
| 0iF2 | 0 | + | 47282308 | 23905822 | 6.18 | | | |
| 0iF3 | 0 | + | 54230058 | 25344368 | 6.02 | | | |
| 0iF4 | 0 | + | 40244798 | 18954394 | 6.43 | | | |
| 0iH1 | 0 | + | 64726838 | 26952996 | 5.79 | 70771 | 1.08 | 1.05 |
| 0iH2 | 0 | + | 46838471 | 22246978 | 6.8 | | | |
| 0iH3 | 0 | + | 40909454 | 18903370 | 6.87 | | | |
| 0iH4 | 0 | + | 54067924 | 25119346 | 6.35 | | | |
| 3iF1 | 3 | + | 54065239 | 10886919 | 5.96 | 67391 | 1.16 | 1.06 |
| 3iF2 | 3 | + | 48714606 | 14470686 | 6.55 | | | |
| 3iF3 | 3 | + | 44688328 | 16294810 | 6.71 | | | |
| 3iF4 | 3 | + | 36970695 | 15737834 | 5.81 | | | |
| 3iF5 | 3 | + | 32077041 | 12065542 | 5.3 | | | |

| | | | | | | | | |
|---|---|---|---|---|---|---|---|---|
| 3iH1 | 3 | + | 46903548 | 9057427 | 7.19 | 62579 | 1.12 | 1.00 |
| 3iH2 | 3 | + | 41743618 | 15654587 | 6.79 | | | |
| 3iH3 | 3 | + | 38888634 | 12484017 | 6.85 | | | |
| 3iH4 | 3 | + | 45598782 | 16242466 | 6.54 | | | |
| 8iF1 | 8 | + | 50536148 | 13430112 | 6.78 | 61139 | 1.26 | 1.05 |
| 8iF2 | 8 | + | 54337061 | 12648440 | 7.11 | | | |
| 8iF3 | 8 | + | 52282170 | 11852948 | 5.21 | | | |
| 8iF4 | 8 | + | 42891267 | 12120423 | 5.43 | | | |
| 8iH1 | 8 | + | 42290670 | 11796387 | 6.72 | 65043 | 1.57 | 1.10 |
| 8iH2 | 8 | + | 39596909 | 14998918 | 5.20 | | | |
| 8iH3 | 8 | + | 39883594 | 12563437 | 5.27 | | | |
| 8iH4 | 8 | + | 28249292 | 8393601 | 4.57 | | | |
| 8iH5 | 8 | + | 43953185 | 10799451 | 7.85 | | | |
| 12iF1 | 12 | + | 39257829 | 13879417 | 7.14 | 70505 | 1.05 | 1.01 |
| 12iF2 | 12 | + | 42214218 | 16259721 | 7.39 | | | |
| 12iF3 | 12 | + | 40921039 | 10536561 | 7.64 | | | |
| 12iF4 | 12 | + | 45775901 | 12512902 | 7.54 | | | |
| 12iF5 | 12 | + | 36779064 | 10269191 | 7.39 | | | |
| 12iH1 | 12 | + | 43772456 | 18806981 | 6.49 | 65492 | 1.14 | 1.01 |
| 12iH2 | 12 | + | 37690946 | 15270494 | 6.95 | | | |
| 12iH3 | 12 | + | 42788391 | 11435578 | 7.54 | | | |
| 12iH4 | 12 | + | 42138119 | 15016298 | 6.48 | | | |

**Supplementary Table 1. ATAC-seq library statistics.** A total of 71 ATAC-seq libraries were generated, with between 3 and 5 biological replicates per treatment. HPA refers to hours post-amputation. + in the iCRT column indicates regenerating animals were pre-incubated in 5µM iCRT14 for two hours prior to amputation and then left in the iCRT14 solution until tissue was collected for library preparation; - in the iCRT column indicates animals were left untreated. Total read pairs refers to the number of raw read pairs generated for each library. Final Mapped Read Pairs refers to the number of read pairs remaining after mitochondrial, duplicated, and unmapped or ambiguously mapped reads were removed. TSS enrichment refers to the fold enrichment in ATAC-seq signal at the transcription start sites of 2000 highly expressed genes relative to regions ± 1kb from the transcription start site. Reproducible Peaks refers to the number of peaks within each treatment group that were found to be biologically reproducible based on at least three pairwise comparisons using an irreproducible discovery rate cutoff of 0.1. The Self-Consistency Ratio refers to the largest fold difference in the number of reproducible peaks recovered from pseudo-replicates (see methods) when comparing biological replicates within a single treatment group. The Rescue Ratio refers to the fold difference in the number of reproducible peaks recovered from a dataset with perfect reproducibility (generated by pooling and then randomly splitting reads from a given treatment group) compared to the number of true biologically reproducible peaks.

**Supplementary Table 2**

| Sample ID | Hpa | iCRT | Total Reads | Final Mapped Reads |
|---|---|---|---|---|
| 0F1 | 0 | - | 29284004 | 21541719 |
| 0F2 | 0 | - | 24597360 | 17966439 |
| 0F3 | 0 | - | 23926341 | 17350620 |
| 0H1 | 0 | - | 25833101 | 18858726 |
| 0H2 | 0 | - | 27241258 | 19701074 |
| 0H3 | 0 | - | 26803090 | 19487287 |
| 3F1 | 3 | - | 24824468 | 18479637 |
| 3F2 | 3 | - | 23840505 | 17694124 |
| 3F3 | 3 | - | 23900532 | 17743737 |
| 3H1 | 3 | - | 25874570 | 19207371 |
| 3H2 | 3 | - | 24172310 | 17898155 |
| 3H3 | 3 | - | 23397937 | 17670977 |
| 8F1 | 8 | - | 23040344 | 17160851 |
| 8F2 | 8 | - | 26019557 | 19216264 |
| 8F3 | 8 | - | 29378123 | 22033550 |
| 8H1 | 8 | - | 25548801 | 19003624 |
| 8H2 | 8 | - | 21601150 | 16154188 |
| 8H3 | 8 | - | 23904011 | 17656019 |
| 12F1 | 12 | - | 26995204 | 18138493 |
| 12F2 | 12 | - | 23703527 | 15445837 |
| 12F3 | 12 | - | 25488245 | 16491319 |
| 12H1 | 12 | - | 34878154 | 22910100 |
| 12H2 | 12 | - | 31662571 | 21545505 |
| 12H3 | 12 | - | 28192288 | 18744249 |
| 0iF1 | 0 | + | 26635485 | 17851347 |
| 0iF2 | 0 | + | 28052169 | 18904815 |
| 0iF3 | 0 | + | 30718370 | 20708864 |
| 0iH1 | 0 | + | 36328076 | 23784082 |
| 0iH2 | 0 | + | 26044439 | 17240456 |
| 0iH3 | 0 | + | 31384073 | 20624671 |
| 8iF1 | 8 | + | 37979732 | 24369941 |
| 8iF2 | 8 | + | 31722061 | 20432162 |
| 8iF3 | 8 | + | 38252155 | 25503172 |
| 8iH1 | 8 | + | 34522120 | 22339316 |
| 8iH2 | 8 | + | 34442705 | 21537194 |
| 8iH3 | 8 | + | 29727022 | 19082280 |
| 12iF1 | 12 | + | 22354289 | 13460197 |
| 12iF2 | 12 | + | 35141637 | 21948916 |
| 12iF3 | 12 | + | 35240879 | 22241372 |
| 12iH1 | 12 | + | 25883305 | 16660844 |
| 12iH2 | 12 | + | 32237790 | 20843729 |
| 12iH3 | 12 | + | 73113728 | 44132834 |

**Supplementary Table 2. RNA-seq library statistics.** A total of 42 RNA-seq libraries were generated, with 3 biological replicates per treatment. HPA refers to hours post-amputation. + in the iCRT column indicates regenerating animals were pre-incubated in 5μM iCRT14 for two hours prior to amputation and then left in the iCRT14 solution until tissue was collected for library preparation; - in the iCRT column indicates animals were left untreated. Total Reads refers to the number of raw reads generated for each library. Final Mapped Reads refers to the number of reads that were successfully mapped to the *Hydra* 2.0 genome gene models.

**Supplementary Table 3**

| Gene Name | Genome Gene Model ID |
|---|---|
| *axin* | g14938 |
| *brachyury1* | g24952 |
| *budhead* | g24126 |
| *cr3l* | g32585 |
| *creb* | g16491 |
| *dishevelled* | g29781 |
| *distal-less* | g18245 |
| *fos* | g23720 |
| *gremlin-like* | g14229 |
| *jun* | g1450 |
| *naked cuticle* | g30442 |
| *nk-2* | g31954 |
| *notum* | g26902 |
| *prdl-a* | g15226 |
| *sFRP* | g12274 |
| *sp5* | g33422 |
| *tcf* | g27364 |
| *wnt3* | g28064 |
| *wnt7* | g29750 |
| *wnt9/10c* | g33373 |
| *wntless* | g18842 |

**Supplementary Table 3. Genome gene model IDs for in-text gene names.** The *Hydra* 2.0 genome gene model IDs that correspond to the gene names used in this study. The *Hydra* 2.0 genome gene models can be found at arusha.nhgri.nih.gov/hydra/download/?dl=nt.

**Supplementary File 1. Full results tables for all pairwise comparisons performed on ATAC-seq data.** Excel workbook containing the results tables generated by edgeR as individual worksheets. Peaks were included in the analysis if they had at least 10 mapped counts per million in at least three samples. The first sheet in the workbook (“Contrasts”) provides the formulae associated with the names assigned to each comparison in the workbook. The “Contrasts” worksheet also lists which consensus peakset was used for each pairwise comparison: either the peakset generated from only untreated samples (“Untreated”) or the peakset generated from all samples (“Full”). The peakset files (“untreated_consensus_diffbind_labels.bed” and “full_consensus_diffbind_labels.bed” respectively) are provided in the repositories associated with this study.

**Supplementary File 1. Full results tables for all pairwise comparisons performed on RNA-seq data.** Excel workbook containing the results tables generated by edgeR as individual worksheets. Genes were included in the analysis if they had at least 2 mapped counts per million in at least three samples. The first sheet in the worksheet (“Contrasts”) provides the formulae linked with the names given to each comparison in the workbook.
